# Identification of brain metastasis genes and therapeutic evaluation of histone deacetylase inhibitors in a clinically relevant model of breast cancer brain metastasis

**DOI:** 10.1101/296384

**Authors:** Soo-Hyun Kim, Richard P. Redvers, Lap Hing Chi, Xiawei Ling, Andrew J. Lucke, Robert C. Reid, David P. Fairlie, Ana Carolina Baptista Moreno Martin, Robin L. Anderson, Delphine Denoyer, Normand Pouliot

## Abstract

Breast cancer brain metastasis remains largely incurable. While several mouse models have been developed to investigate the genes and mechanisms regulating breast cancer brain metastasis, these models often lack clinical relevance since they require the use of immune-compromised mice and/or are poorly metastatic to brain from the mammary gland. We describe the development and characterisation of an aggressive brain metastatic variant of the 4T1 syngeneic model (4T1Br4) that spontaneously metastasises to multiple organs, but is selectively more metastatic to the brain from the mammary gland than parental 4T1 tumours. By immunohistochemistry, 4T1Br4 tumours and brain metastases display a triple negative phenotype, consistent with the high propensity of this breast cancer subtype to spread to brain. *In vitro* assays indicate that 4T1Br4 cells have an enhanced ability to adhere to or migrate across a brain-derived endothelial monolayer and greater invasive response to brain-derived soluble factors compared to 4T1 cells. These properties are likely to contribute to the brain-selectivity of 4T1Br4 tumours. Expression profiling and gene set enrichment analyses demonstrate the clinical relevance of the 4T1Br4 model at the transcriptomic level. Pathway analyses implicate tumour-intrinsic immune regulation and vascular interactions in successful brain colonisation, revealing potential therapeutic targets. Evaluation of two histone deacetylase inhibitors, SB939 and 1179.4b, shows partial efficacy against 4T1Br4 metastasis to brain and other sites *in vivo* and potent radio-sensitising properties *in vitro*. The 4T1Br4 model provides a clinically relevant tool for mechanistic studies and to evaluate novel therapies against brain metastasis.

**SUMMARY STATEMENT:** We introduce a new syngeneic mouse model of spontaneous breast cancer brain metastasis, demonstrate its phenotypic, functional and transcriptomic relevance to human TNBC brain metastasis and test novel therapies.

## INTRODUCTION

Metastasis accounts for the majority of breast cancer-related mortalities and approximately 40,000 women were expected to die from the disease in 2017 in the US alone (Ghoncheh et al., 2016). The incidence of brain metastasis in breast cancer patients is estimated at ~30% and is rising, despite more effective systemic therapies (Tabouret et al., 2012). Therapeutic options consist primarily of surgery, whole brain radiation therapy, stereotactic radiosurgery and chemotherapy but these approaches, while extending life, are rarely curative (Eichler et al., 2011). While surgical resection improves the overall median survival in the range of 10-16 months, surgery is often not feasible for patients with multiple or large brain metastases and progressive extra-cranial disease. Cognitive impairment due to damage to the surrounding normal tissue also limits the dose and therapeutic efficacy of radiation therapy (Lee et al., 2012; McTyre et al., 2013). Moreover, the efficacy of chemotherapy against brain metastases is limited by the poor penetration of most chemotherapeutic drugs through the blood-brain barrier (BBB) and/or by the acquisition of resistance once brain metastases are established (Arslan et al., 2014; Lockman et al., 2010).

Breast cancer patients with tumours over-expressing human epidermal growth factor receptor 2 (HER2) or triple negative breast cancer (TNBC) that lacks expression of oestrogen (ER) and progesterone receptors (PR) and does not have amplification of HER2, have a disproportionately high incidence of brain recurrence and tend to develop brain metastases early after initial diagnosis (Heitz et al., 2009). While HER2-targeted therapies using Trastuzumab or Lapatinib have been developed, these inhibitors alone or in combination with chemotherapy extend life primarily by providing effective control of extra-cranial disease but have limited efficacy against TNBC brain metastases (Bendell et al., 2003; Burstein et al., 2005; Chen et al., 2013; Liu et al., 2011). Hence, the molecular mechanisms driving TNBC brain metastasis remain poorly understood and requires the identification of new therapeutic targets.

A number of animal models of breast cancer brain metastasis have been developed to investigate these mechanisms and to evaluate new therapies (Daphu et al., 2013). However, most require injection of a large bolus of cells into the left cardiac ventricle or in the carotid artery for efficient dissemination to brain (Bos et al., 2009; Lockman et al., 2010; Lorger and Felding-Habermann, 2010; Palmieri et al., 2007; Simoes et al., 2008). While informative, such experimental metastasis models lack clinical relevance since they bypass the formation of a primary tumour and only recapitulate late stages of the metastatic process. Accordingly, they are not appropriate for investigating the efficacy of preventive or neo-adjuvant therapies against spontaneous brain metastasis. Moreover, the majority of these models are human xenografts that require the use of immune deficient mice and do not faithfully replicate the pro- and anti-tumour properties of the immune system (Hamilton and Sibson, 2013).

Transgenic mouse models or syngeneic tumour transplantation models, in which murine tumour cells are inoculated into the mammary fat pad, better mimic metastasis in patients. They recapitulate the entire metastatic process, from formation of a primary tumour to the spontaneous spread of metastatic cells to multiple organs, in immune competent animals (Daphu et al., 2013; Erin et al., 2013). Few of these models, however, are spontaneously metastatic to brain. The 4T1 syngeneic mouse model of metastasis has been reported to spread to brain from the mammary gland, but with a relatively low incidence ranging from 2% to 25% (Bailey-Downs et al., 2014; Tao et al., 2008; Zhang et al., 2015). Recently, we developed an aggressive 4T1-derived brain metastatic tumour variant (4T1Br4) that consistently gives rise to a high incidence of spontaneous brain metastases (Martin et al., 2017). In this model we demonstrated that [10]-gingerol partially reduced spontaneous brain metastasis (Martin et al., 2017). While 4T1-derived spontaneous metastasis models represent a significant advance over xenograft and experimental models of brain metastasis, they require thorough validation of their clinical and molecular relevance to human breast cancer brain metastasis (Hollern and Andrechek, 2014). Here we describe the development of the 4T1Br4 spontaneous breast cancer brain metastasis model in more detail and demonstrate its phenotypic, functional and transcriptomic relevance to human brain-metastatic TNBC. We identify new brain metastasis genes and signalling pathways likely to contribute to the spread and development of TNBC brain lesions. Preliminary evaluation of the efficacy of two novel histone deacetylase inhibitors (HDACi) indicates partial efficacy against spontaneous brain metastasis and demonstrate that the 4T1Br4 model is well suited for profiling novel therapeutics against brain-metastatic TNBC.

## RESULTS

### Development of the 4T1Br4 model of spontaneous brain metastasis

The 4T1 syngeneic mouse model has been widely used to investigate genes and mechanisms regulating breast cancer metastasis. Tumours derived from the parental 4T1 tumour line spontaneously spread to multiple organs following their inoculation in the mammary fat pad and give rise to extensive lung metastases, moderate level of bone metastases but only low incidence of brain metastases (Bailey-Downs et al., 2014; Kusuma et al., 2012; Tao et al., 2008; Zhang et al., 2015). A schematic of the procedure to develop a more robust model of spontaneous brain metastasis is presented in Supplementary Material Fig. S1A. Briefly, parental 4T1 cells were tagged with a mCherry fluorescent marker gene for visualising brain lesions at endpoint and quantitating metastatic burden by genomic qPCR (Martin et al., 2017). The cells were subjected to four rounds of *in vivo* enrichment by inoculation of cells in the mammary gland of BALB/C mice, followed by explant cultures of excised brain lesions, fluorescence-activated cell sorting (FACS) and *in vitro* expansion of mCherry^+ve^ cells to generate the 4T1Br4 line. At this stage the incidence of spontaneous brain metastasis reached 20% in mice bearing bulk 4T1Br4 tumours compared to approximately 7% in mice bearing parental 4T1 tumours. Clonal variants were then isolated from the bulk population of 4T1Br4 cells by FACS and individually tested *in vivo* using a low cell number (2 × 10^4^/mouse) to avoid rapid metastatic progression to other organs and/or excessive tumour size requiring early termination of the experiment (Bailey-Downs et al., 2014). Under these conditions, we found extensive variability in the incidence of brain metastasis between clones ranging from 0% to 80% (despite similar primary tumour growth rate), indicating that some heterogeneity remains even after serial *in vivo* selection (Supplementary material Fig. S1B). Clone 6 (67% incidence) was selected for subsequent experiments and is referred to as 4T1Br4 for brevity.

The growth rate of 4T1 and 4T1Br4 orthotopic tumours did not differ significantly, indicating that increased brain metastasis in the 4T1Br4 model is not due to faster tumour formation *in vivo* (Fig. 1A). In the next series of experiments, tumours were surgically resected when they reached ~0.4 - 0.5 g to better reflect the clinical situation and to allow more time for late brain metastases to develop while avoiding growth of primary tumours to unethical size (Fig. 1B). In this setting, 4T1Br4 tumours were metastatic to multiple organs, but were selectively more metastatic to brain compared to parental 4T1 tumours (Fig. 1C-E).

**Fig. 1.**
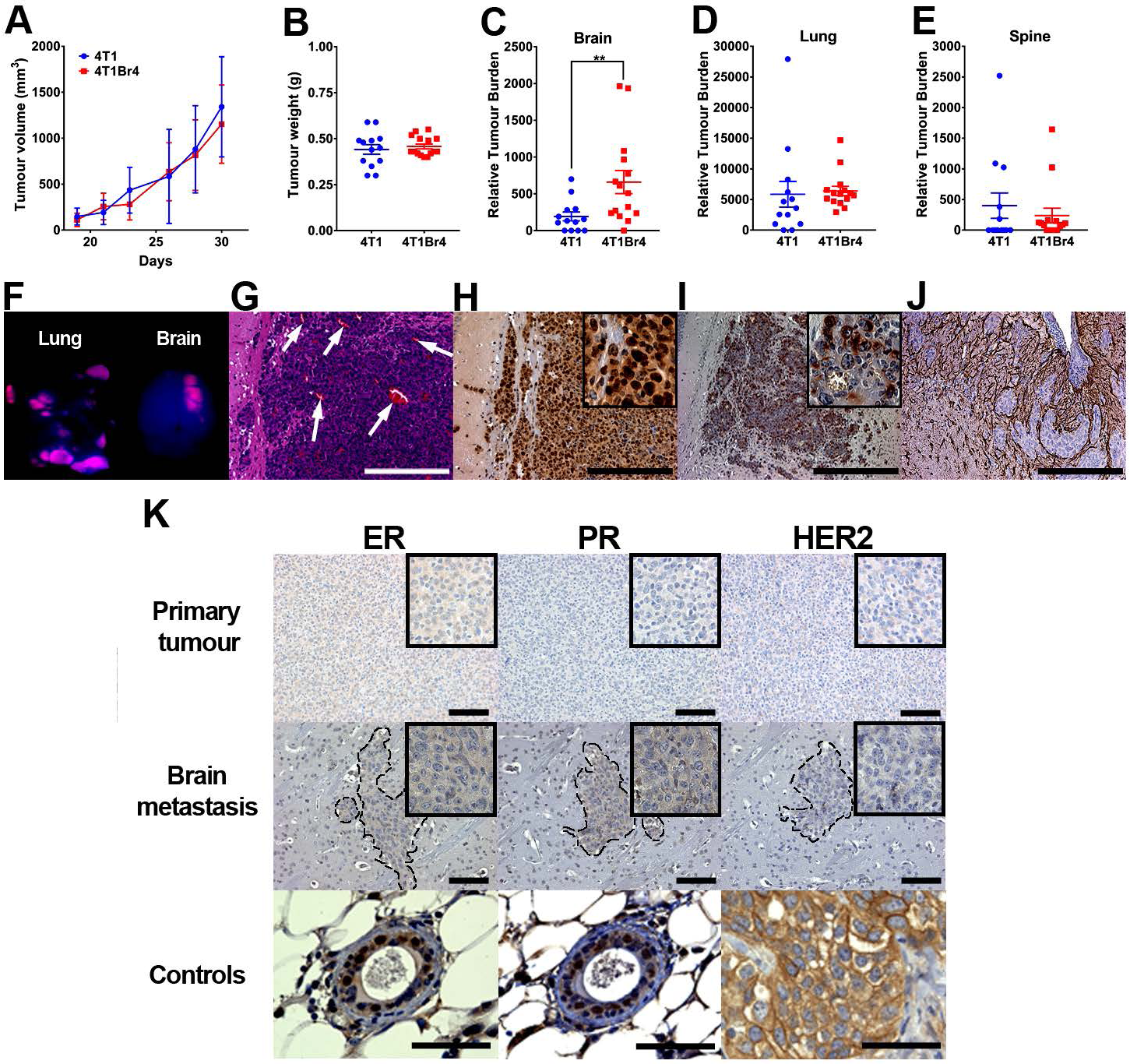
Metastatic and phenotypic characterisation of the 4T1Br4 model. (A) Mice were inoculated orthotopically with 4T1Br4 cells (2 × 10^4^) and tumour growth measured between day 19 and day 30. Data show mean ± SD from 15 mice per group. (B) For metastasis assays, 4T1 (n = 13) and 4T1Br4 (n = 15) primary tumours were resected when they reached approximately 0.4 - 0.5 g. Relative tumour burden in (C) brain, (D) lung and (E) spine was measured 3 weeks after tumour resection. Each dot in (B-D) represents one mouse and the horizontal line represents mean ± SEM. **p<0.01 (Student’s t-test). (F) Fluorescence imaging of mCherry^+ve^ (pink) cells in lung and brain metastases. (G) H&E staining of a brain macrometastasis (arrows, blood vessels). (H) Ki-67, (I) pan-cytokeratin and (J) GFAP were detected by standard immunohistochemistry. Scale bar = 100 μm. High power images of (H) and (I) are shown in insets. (K) IHC detection of ER, PR and HER2 expression in 4T1Br4 primary tumours (top panels) and brain metastases (middle panels). Metastatic lesions are delineated by a dotted line. Hematoxylin was used for nuclear staining (blue). High power images are shown in insets. Normal mammary glands were used as positive controls for nuclear expression of ER and PR and a human SKBR3 primary tumour xenograft was used as a positive control for membrane expression of HER2 (bottom panels). Scale bar = 50 μm.

*Ex-vivo* fluorescence imaging of organs from 4T1Br4-bearing mice revealed the presence of multiple lesions in bone and soft tissues including lung and brain. In contrast to the lungs in which multiple lesions were evident, mice typically developed a single brain macrometastasis, most commonly observed in the cerebral cortex (Fig. 1F) but also detected occasionally in the cerebellum and leptomeninges (not shown). Brain metastases were highly vascularised (Fig. 1G), proliferative as evidenced by Ki67 IHC staining (Fig. 1H), expressed cytokeratins (Fig. 1I) and were always surrounded by glial fibrillary acidic protein (GFAP^+ve^) activated astrocytes (Fig. 1J), consistent with a reactive glia (Fitzgerald et al., 2008).

The triple negative status of 4T1Br4 primary tumours was confirmed by IHC staining of ER, PR and HER2. As expected, 4T1Br4 tumours showed no evidence of nuclear ER and PR expression and lacked cell surface expression of HER2 (Fig. 1K, top panels). Discordance in receptor status between primary tumours and brain metastases has been reported in some patients (Duchnowska et al., 2012). This phenotypic receptor conversion could potentially lead to inappropriate treatment and lack of response to therapy in patients with advanced disease. Hence, we further evaluated ER, PR and HER2 expression in brain metastases. Only weak cytoplasmic (but not nuclear) staining for ER and PR was observed occasionally in some lesions and no HER2 expression was detected indicating that 4T1Br4 brain metastases maintain their triple negative status (Fig. 1K, middle panels).

### 4T1Br4 cells demonstrate enhanced adhesive, migratory and invasive abilities *in vitro* that are likely to contribute to brain metastasis *in vivo*

Surrogate *in vitro* assays were used to gain further insights into the potential functional attributes that contribute to 4T1Br4 preferential metastasis to the brain. Parental 4T1 and 4T1Br4 cell showed no significant differences in proliferative rate (Fig. 2A) or in their ability to form colonies at low density (Fig. 2B). In contrast, 4T1Br4 cells were more migratory than 4T1 cells in serum chemotaxis assays (Fig. 2C). Importantly, 4T1Br4 cells were significantly more adherent to (Fig. 2D) and migrated more efficiently than parental 4T1 cells through a monolayer of brain microvascular endothelial cells (Fig. 2E).

**Fig. 2.**
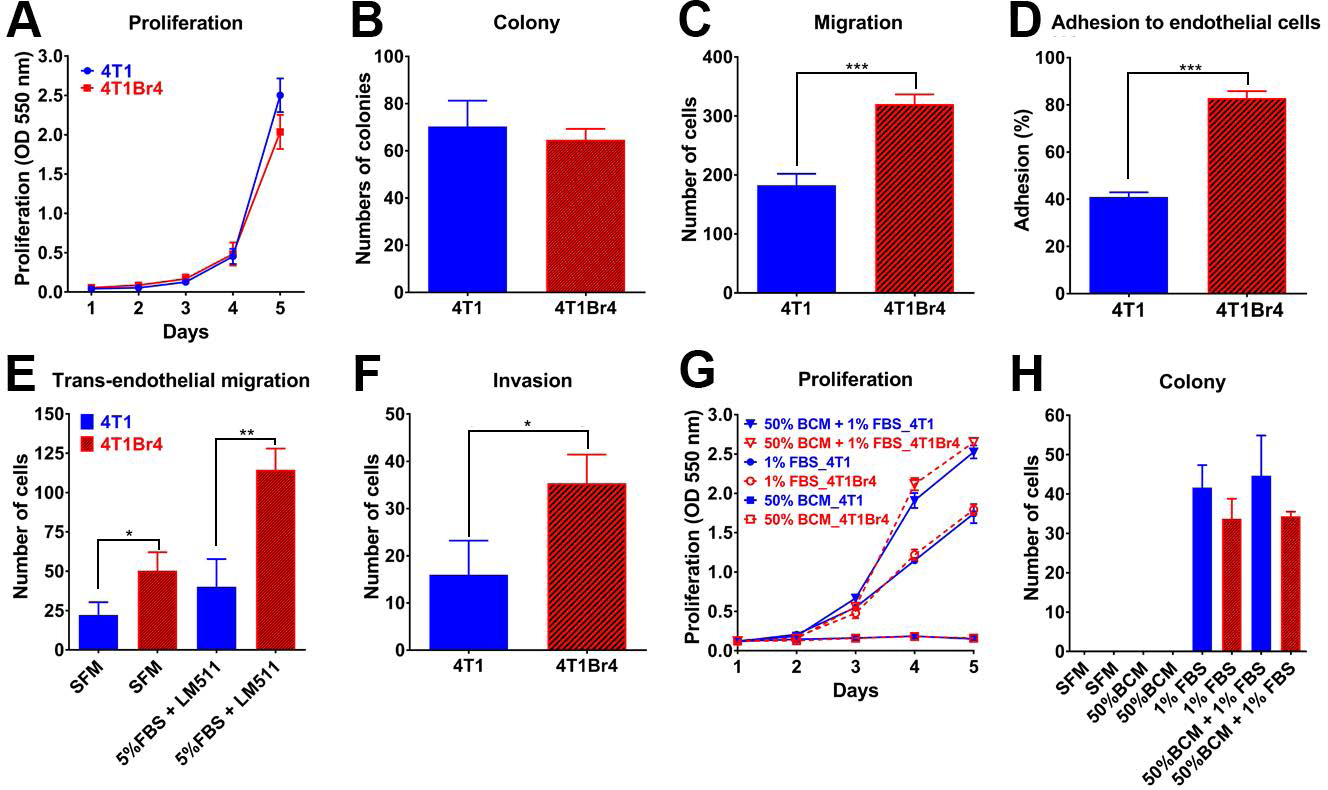
Functional characterisation of 4T1 and 4T1Br4 cell *in vitro*. (A) Proliferation assay. Cells were seeded at 500 cells/well in 96-well plates in the presence of 5% FBS and cultured for 5 days. (B) Colony forming assay. Cells were seeded at low density (100 cells/well in 6-well plates) in the presence of 5% FBS and colonies were counted after 10 days. (C) Migration assay. Cells were seeded in the upper well of Transwell chambers in serum-free medium and 5% FBS added to the lower well as chemoattractant. Cells that migrated to the underside of the porous membrane were counted after 4 hours at 37°C. (D) Adhesion assay. Calcein-labelled cells were seeded onto a monolayer of bEnd.3 brain microvascular endothelial cells and tumour cells attached to bEnd.3 cells were counted as described in Materials & Methods. (E) Trans-endothelial migration assay. Tumour cells were seeded onto a monolayer of bEnd.3 cells in the upper well of Transwell chambers and tumour cells that migrated to the underside of the membrane in response to serum-free medium (SFM) or 5% FBS + laminin-511 in the lower well were counted after 48 hours incubation at 37°C. (F) Invasion assay. Tumour cells were mixed with Matrigel (1:1 ratio) in the upper well of Transwell chambers and cells that invaded to the underside of the membrane in response to 50% brain conditioned medium (BCM) in the lower well were counted 18 hours after cell seeding. (G) Proliferation and (H) colony forming assays. Cells were seeded in the presence of 50% BCM, 1% FBS or both as indicated and cultured for 5 days (G) or 10 days (H). All experiments were conducted in triplicate and repeated three times (n = 3). Data show mean ± SD of triplicates from a representative experiment (n = 3). p-values were calculated using a two-ANOVA (A, G), Mann-Whitney non-parametric test (B, C, D, F) or one-way ANOVA, Bonferroni post-test (E, H); p<0.05 was considered significant. *p < 0.05, **p < 0.01, ***p < 0.005.

For invasion assays, serum-free brain conditioned medium (BCM) was prepared from primary neonatal whole brain explant cultures and used as a chemoattractant. 4T1Br4 were significantly more invasive than 4T1 cells in response to 50% BCM (Fig. 2F). In contrast, BCM was insufficient alone to promote 4T1Br4 or 4T1 cell proliferation (Fig. 2G). Whilst BCM enhanced proliferation in combination with low serum concentrations (1% v/v), it did so to the same extent in 4T1 and 4T1Br4 cells (Fig. 2G) indicating that proliferation in response to brain-derived growth factors is unlikely to be a major factor contributing to the brain selectivity of 4T1Br4 tumours *in vivo*. Similarly, BCM alone did not promote colony formation, nor did it enhance colony formation under low serum conditions in either 4T1 or 4T1Br4 cells (Fig. 2H). Collectively, these results indicate that 4T1Br4 cells have acquired several functional properties likely to contribute to their greater brain-metastatic abilities, most notably increased endothelial adhesion, trans-endothelial migration and invasive response to brain-derived soluble factors.

### Evaluation of HDAC inhibitor efficacy in the 4T1Br4 model

There is increasing interest in the use of HDACi for cancer therapy (Li and Seto, 2016). HDACs regulate multiple processes essential for cancer progression including cell cycle, differentiation, cell motility, autophagy, apoptosis and angiogenesis, by de-acetylating histones and non-histone proteins and altering gene transcription. Changes in their expression and/or activity are common in tumours (Nakagawa et al., 2007; Weichert, 2009). Importantly, SAHA (Vorinostat), a pan-HDAC inhibitor (Richon et al., 2009) has been shown previously to cross the BBB and to partially inhibit brain colonisation in the MDA-MB-231Br experimental metastasis xenograft model (Palmieri et al., 2009).

To demonstrate the clinical relevance of the 4T1Br4 model for therapy testing, we assessed the efficacy of two novel HDACi, SB939 (Pracinostat) and 1179.4b (Fig. 3A). Both compounds have shown selectivity towards tumour cells compared to primary fibroblasts *in vitro* and higher potency than SAHA against human breast tumour cells (Kahnberg et al., 2006; Novotny-Diermayr et al., 2010). However, their efficacy has not been evaluated in pre-clinical models of breast cancer brain metastasis. First, we compared the anti-tumour effect of SB939 and 1179.4b to that of SAHA *in vitro*. In colony formation assays, SAHA (1 μM) inhibited 4T1Br4 colony formation by approximately 30%. By comparison, SB939 and 1179.4b inhibited colony formation by ~50% and 100%, respectively, at the same concentration (Fig. 3B). The effect of SB939 was evident from 500 nM whereas 1179.4b partially inhibited colony formation at concentration as low as 31.25 nM (~30%) and completely blocked colony formation at 250 nM or above (Fig. 3C). The IC_50_ values of these compounds were determined using a 3-day proliferation assay. SB939 and 1179.4b showed greater inhibitory potency than SAHA against 4T1Br4 (Fig. 3D) and MDA-MB-231Br (Fig. 3E). IC_50_ values for SAHA were estimated at 1.3 µM and 1.7 µM in 4T1Br4 and MDA-MB-231Br cells respectively (Fig. 3F), in good agreement with IC_50_ values reported in other tumour models (Atadja, 2009). In contrast, IC_50_ for SB939 were 527 nM and 364 nM and for 1179.4b, 71 nM and 50 nM in 4T1Br4 and MDA-MB-231Br respectively. Taken together, these results indicate that SB939 and 1179.4b are significantly more potent than SAHA in inhibiting brain-metastatic breast tumour cells *in vitro*.

**Fig. 3.**
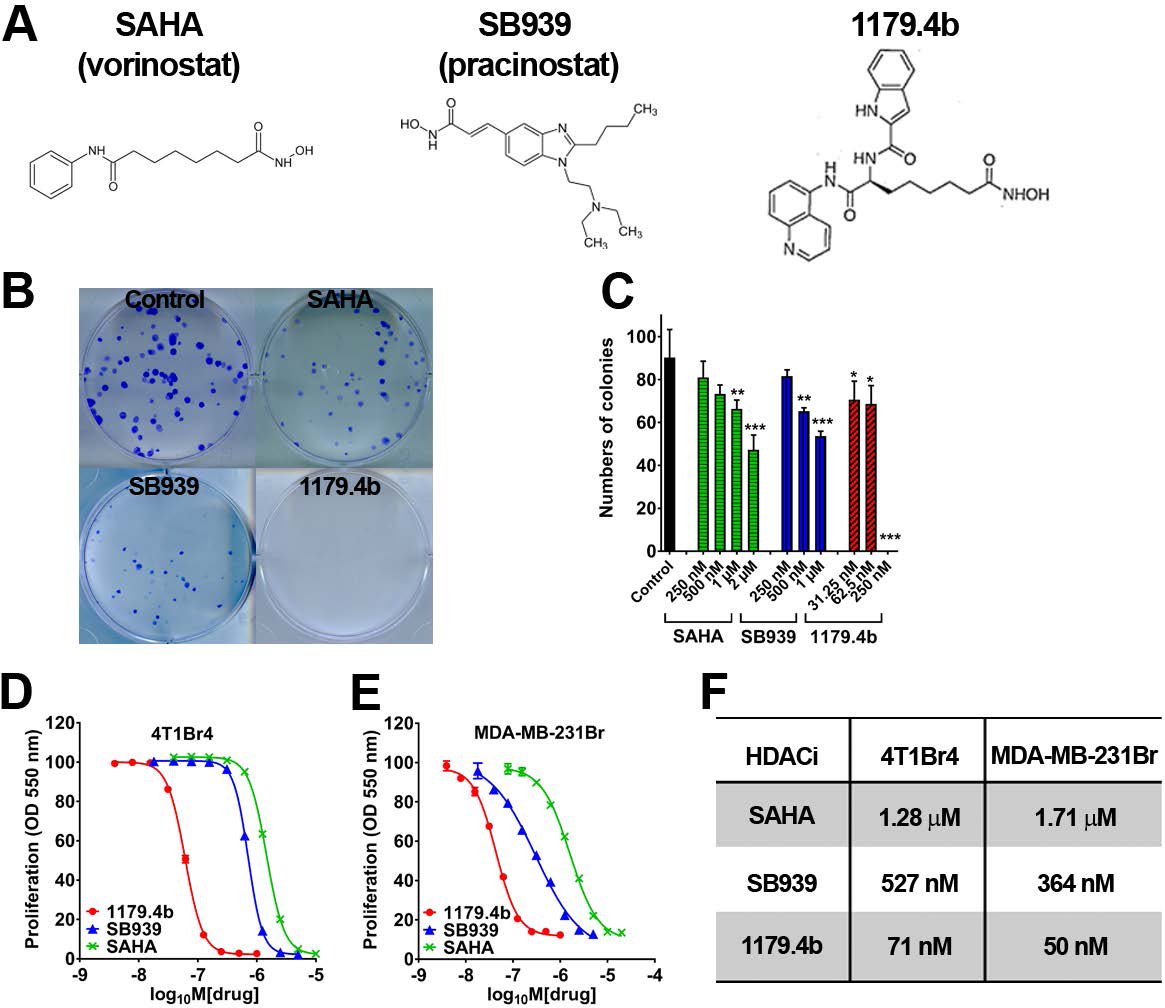
Cytotoxic effect of HDACi on brain-metastatic breast tumour cells *in vitro*. (A) Molecular structure of SAHA, SB939 and 1179.4b. (B) Colony formation assay. 4T1Br4 cells (1 × 10^2^/well) were seeded in 6 well plates, allowed to attach for 6 hours at 37°C and adherent cells treated with 1 μM of SB939 or 1179.4b or SAHA or DMSO (vehicle control) as indicated. Colonies (> 50 cells) formed after 10 days were stained with crystal violet and counted. Representative images of colonies from 3 independent experiments performed in triplicates are shown. (C) Colony formation was assayed as in (B) at the indicated concentrations of HDACi. Data show average number of colonies/culture condition ± SD of triplicate wells from a representative experiment (*p < 0.05, **p < 0.01, ***p<0.001 compared to DMSO control). p-values were calculated using a one-way ANOVA with Bonferroni post-test; p < 0.05 was considered significant. (D) 4T1Br4 (1 × 10^3^/well) and (E) MDA-MB-231Br cells (2 × 10^3^/well) were seeded in 96-well plates and treated with increasing doses of SB939 (blue), 1179.4b (red) or SAHA (green) for 72 hours at 37°C. Proliferation was measured by colorimetric SBR assay. Data show a representative experiment (n = 3) and are expressed as means ± SEM of six replicate wells per condition. (F) IC_50_ values for SAHA, SB939 and 1179.4b were derived from (D) and (E) for 4T1Br4 and MDA-MB-231 Br cells respectively and calculated using Hill’s equation in the GraphPad Prism 6.0 software.

Acetylation of histone H3 is associated with transcriptional activation of several genes involved in the suppression of tumour growth and was suggested to predict clinical outcome in cancer patients (Novotny-Diermayr et al., 2011; Seligson et al., 2009). To determine if SB939 and 1179.4b induce hyperacetylation of histone H3 in brain-metastatic lines, 4T1Br4 and MDA-MB-231Br cells were examined by immunoblotting following exposure to SB939 or 1179.4b for 24 hours. Histone H3 acetylation increased sharply in 4T1Br4 cells in response to SB939 at concentrations ≥ 1.2 µM whereas treatment with 1179.4b induced hyperacetylation between 312 nM and 5 µM (Supplementary material Fig. S2A). In MDA-MB-231Br cells, SB939 induced a dose-dependent response between 625 nM and 5 µM, while histone H3 hyperacetylation in response to 1179.4b was evident at all concentrations tested (312 nM - 5 µM) (Fig. S2B). Histone H3 hyperacetylation was maximal at 24 hours for both compounds in both cell lines, albeit with a slower kinetic in 4T1Br4 cells than in MDA-MB-231Br cells (data not shown).

To assess whether hyperacetylation of histone H3 in response to SB939 or 1179.4b is reversible, cells were treated with each HDACi for 24 hours and changes in the level of histone H3 acetylation in 4T1Br4 (Supplementary material Fig. S2C) and MDA-MB-231Br cells (Supplementary material Fig. S2D) were measured over 24 hours after removal of the inhibitors. 4T1Br4 cells treated with SB939 (2 μM) showed a rapid decrease in the level of histone H3 acetylation following drug removal whereas acetylation remained detectable for up to 8 hours in MDA-MB-231Br cells. Histone H3 acetylation in response to 1179.4b (1 μM) was detectable for up to 4 hours in 4T1Br4 cells and up to 24 hours in MDA-MB-231Br cells. Collectively, these data demonstrate that both SB939 and 1179.4b reversibly induce histone H3 hyperacetylation in mouse and human brain metastatic breast cancer cell lines and that 1179.4b sustains acetylation of histone H3 longer in both cell lines compared to SB939. These observations are consistent with their respective potency demonstrated in colony formation and proliferation assays.

Neither SB939 nor 1179.4b has been evaluated in breast cancer brain metastasis models. In preliminary experiments we confirmed that SB939 or 1179.4b (both at 10, 20 or 40 mg/kg) administered intraperitoneally once daily for 8 days in BALB/C mice were well tolerated, with no significant decrease in body weight or changes in overall appearance or behaviour (data not shown). However, 1179.4b (40 mg/kg) had poor solubility in 30% polyethylene glycol, and therefore a dose of 20 mg/kg was chosen for subsequent *in vivo* experiments. For SB939, the dose was increased to 50 mg/kg since it retained good solubility at this concentration and was well tolerated at the highest dose tested (40 mg/kg). To investigate the effect of SB939 and 1179.4b on tumour growth *in vivo*, 4T1Br4 cells were injected into the mammary fat pad and drug treatment initiated on day 9, when primary tumours were palpable. SB939 (50 mg/kg) and 1179.4b (20 mg/kg) were injected intraperitoneally once daily until completion of the experiment on day 28. Both SB939 and 1179.4b significantly reduced 4T1Br4 primary tumour growth (Fig. 4A) and tumour weight at endpoint (Fig. 4B) compared to control mice. Primary tumours were analysed for changes in histone H3 acetylation by immunoblotting (Fig. 4C). As expected, SB939- or 1179.4b-treated mice showed a strong increase in acetylation of histone H3 in primary tumours compared to tumours from control mice, confirming that both SB939 and 1179.4b effectively blocked HDAC activity *in vivo* and caused reduced primary tumour growth.

**Fig. 4.**
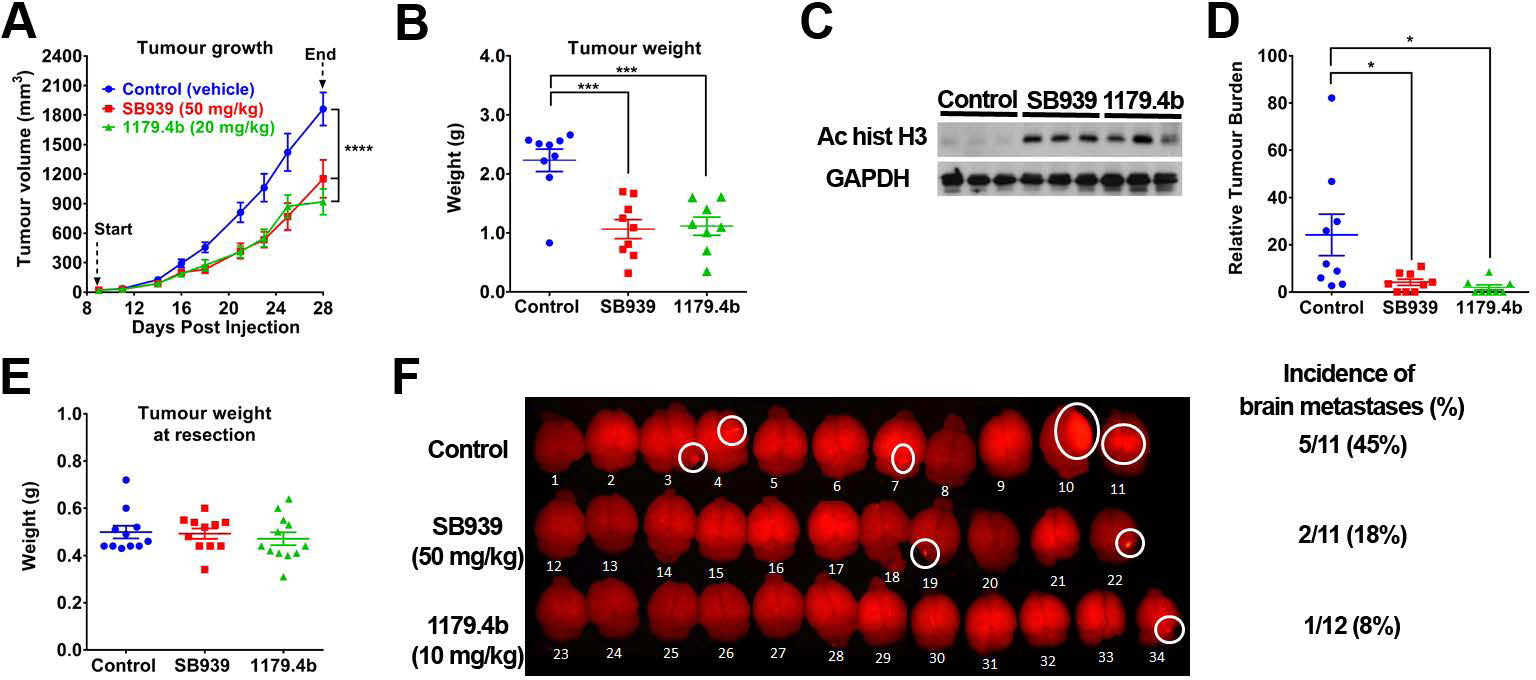
SB939 and 1179.4b inhibit 4T1Br4 tumour growth and spontaneous metastasis. (A) 4T1Br4 (2 × 10^4^) cells were injected into the mammary fat pad of mice and tumour growth measured thrice weekly with electronic callipers. SB939 (50 mg/kg, red), 1179.4b (20 mg/kg, green) and vehicle (control) were administered intraperitoneally once daily from day 9 to day 28 and the mice harvested one hour after the last drug treatment. (B) 4T1Br4 primary tumours were collected and weighed. (C) Whole tumour lysates of three representative tumours from each experimental group were analysed for acetylation of histone H3 by western blotting. GAPDH was used as loading control. (D) Metastatic burden in spines was analysed by genomic qPCR of the mCherry marker gene. Data in (A, B and D) are expressed as means ± SEM of tumour volume/tumour weight/metastatic burden in spine, with each dot in (B) and (D) representing one mouse. Control (n = 9), SB939 (n = 9) and 1179.4b (n = 8). p-values were calculated using a two-way ANOVA with a Tukey post-test for tumour growth, one-way ANOVA with a Bonferroni post-test for tumour weight and metastatic burden in spines. *p < 0.05, ***p < 0.01, ****p < 0.001 compared to control (vehicle). (E) For spontaneous brain metastasis, primary tumours were resected when they reached ~ 0.5 g and the mice allocated to control (n = 11), SB939 (n = 11) or 1179.4b (n = 12) treatment group. Data show means ± SEM, with each dot representing one mouse. (F) Two days after tumour resection, mice were treated once daily with SB939 (50 mg/kg) or 1179.4b (10 mg/kg) by intraperitoneal injection and brains were imaged *ex-vivo* by fluorescence for detection of mCherry^+ve^ lesions at endpoint. Incidence of detectable brain metastases is shown on the right.

While the primary objective of this experiment was to evaluate the impact of SB939 and 1179.4b on primary tumour growth, spines were also collected and relative tumour burden was measured by genomic qPCR detection of the mCherry marker gene. Interestingly, metastatic burden in spines was significantly reduced by SB939 or 1179.4b (Fig. 4D). Brain metastasis was not evaluated in this experiment since fewer mice/group (8-9) were used, tumours were not resected, and the experiment had to be terminated earlier (day 28) due to the large primary tumour size/weight in the control group and early signs of lymphopenia occurring in mice treated for extended time (19 days) with 1179.4b.

To investigate the effects of SB939 and 1179.4b specifically on spontaneous brain metastasis, 4T1Br4 cells were injected into the mammary fat pad and the tumours resected when they reached approximately 0.5 g. Treatment with SB939 (50 mg/kg) or 1179.4b (10 mg/kg) was initiated two days after tumour resection. Tumour weights at resection were not significantly different between groups (Fig. 4E). Brains were examined by fluorescence imaging at endpoint (day 33) and the incidence of mice with detectable mCherry^+ve^ lesions was scored (Fig. 4F). Under those conditions, SB939 and 1179.4b reduced the incidence of mice with detectable brain lesions from 45% (5/11) in the control group to 18% (2/11) and 8% (1/12) in SB939 and 1179.4b-treated mice, respectively.

The results above indicate that SB939 and 1179.4b inhibit spontaneous brain metastasis but do not address whether they do so by preventing the establishment of brain metastases or by inhibiting directly their outgrowth in the brain. We addressed this using an experimental metastasis assay in which tumour cells are inoculated directly into the vasculature, with treatment commencing 2 days after cell inoculation and continuing until day 12. The presence of brain metastases at endpoint was confirmed by scoring the extent of cytokeratin^+ve^ tumour cells by IHC in whole brain sections. Experimental brain metastasis was inhibited partially by 1179.4b, but not by SB939 in this protocol (Fig. 5A). The presence of viable circulating tumour cells (CTC) in blood collected from each experimental group was evaluated in a colony formation assay. Consistent with the inhibition of metastasis to brain and other soft tissues observed in 1179.4b-treated animals, the number of colonies formed was dramatically reduced compared to control, whereas SB939 had no significant effect (Fig. 5B). These results indicate that 1179.4b may inhibit brain metastasis primarily by targeting CTCs, thereby preventing their homing and outgrowth in brain (and other organs) rather than inhibiting established metastases directly. Consistent with this possibility, metastases in other soft tissue were also scored semi-quantitatively by visual examination at necropsy and showed a reduction in the incidence of ovary, kidney and adrenal gland metastases in 1179.4b- but not SB939-treated mice (Fig. 5C).

**Fig. 5.**
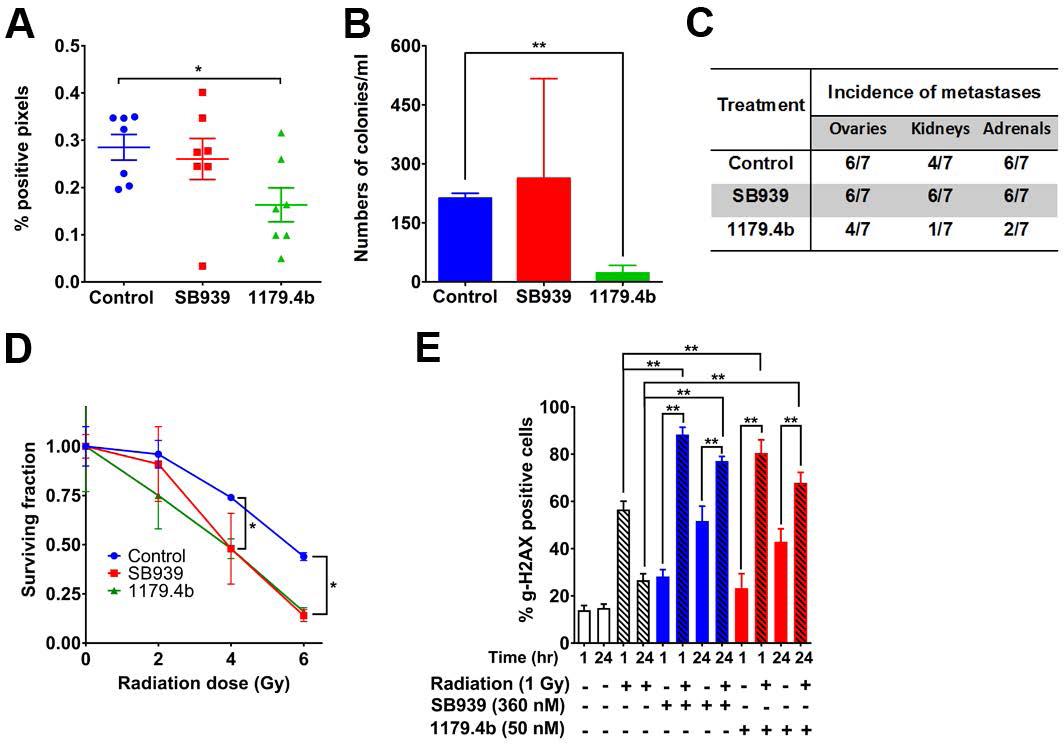
Effect of SB939 and 1179.4b on experimental brain metastasis and radio-sensitivity. (A) 4T1Br4 cells (5 × 10^4^) were inoculated directly into the left cardiac ventricle and mice were administered vehicle (control), SB939 (50 mg/kg) or 1179.4b (20 mg/kg) daily starting 2 days after cell inoculation. Metastasis was scored on day 12 by IHC staining of cytokeratin^+ve^ cells and quantitated using Aperio ImageScope. Data are expressed as % of positive pixels in whole brain sections and show mean ± SEM of 3 step sections/brain, 200 µm apart, with each dot representing one mouse (n = 7 mice/group). (B) Blood was collected by cardiac puncture at endpoint and the number of viable CTCs determined by colony formation assay. Data show mean ± SEM from 4 mice/group. (C) Incidence of experimental metastasis in soft tissues was estimated by visual inspection at necropsy. (D) Adherent 4T1Br4 cells were treated with SB939 (530 nM, red) or 1179.4b (70 nM, green) or vehicle alone (blue) for 24 hours followed by irradiation at the indicated doses. The plates were incubated for 10 days and colonies (> 50 cells) were counted. Data show a representative experiment (n = 3) and expressed as mean surviving fraction ± SD of triplicate wells. (E) MDA-MB-231Br cells (4 × 10^4^/well) were seeded in chamber slides, allowed to adhere for 18 hours at 37°C and treated with SB939 (360 nM) or 1179.4b (50 nM) for 24 hours. Cells were irradiated with 1 Gy and incubated for an additional 1 hour or 24 hours at 37°C before analysis of γ-H2AX foci formation by immunofluorescence. Data show a representative experiment (n = 3) and are expressed as means ± SEM of six fields of view (2 replicate wells × 3 fields of view per condition). p-values in (D) and (E) were calculated using a one-way ANOVA with Bonferroni post-test. *p < 0.05, **p < 0.01.

### SB939 and 1179.4b sensitise brain-metastatic cells to radiation

Whole brain radiation therapy (WBRT) remains the cornerstone of treatment for brain metastasis and its efficacy is increasingly being explored in combination with other therapeutic modalities (Tallet et al., 2017). Several HDACi, including SAHA, have been reported to sensitise tumour cells to radiation (Baschnagel et al., 2009; Chiu et al., 2013), a strategy that has the potential to increase the efficacy of radiation therapy while minimising its side effects. Therefore, the radiosensitising properties of SB939 and 1179.4b were evaluated first in 4T1Br4 cells using *in vitro* colony forming assays. Each HDACi was used at their IC_50_ concentration either alone or in combination with increasing radiation doses. Both SB939 and 1179.4b significantly enhanced radiation-induced cell death, resulting in fewer colonies compared to radiation alone (Fig. 5D). Dose enhancement factor (DEF) at 50% cell survival was 1.45 for both SB939 and 1179.4b.

Since MDA-MB-231Br cells are highly motile and tend to scatter in colony assays, we used an alternative method to quantitate the radio-sensitising effects of SB939 and 1179.4b by measuring the induction of γ-H2AX nuclear foci. γ-H2AX is recruited to DNA double strand breaks induced by ionising radiation and is commonly used as a marker for DNA damage (Sharma et al., 2012). Sustained γ-H2AX induction (>24 hours) usually indicates inability to repair extensive DNA damage induced by ionising agents and was suggested to be a predictor of tumour radiosensitivity and cytotoxicity (Denoyer et al., 2015; Taneja et al., 2004). A low level of γH2AX was observed in control untreated cells at the 1 hour and 24 hour time-points (Fig. 5E). Radiation alone significantly increased γH2AX foci one hour post-radiation in both cell lines but the level returned to baseline after 24 hours. In contrast, SB939 or 1179.4b alone induced accumulation of γH2AX between 1 and 24 hours, suggesting failure to repair DNA damage. Importantly, combination of either SB939 or 1179.4b with radiation induced a strong γH2AX response (~80% of cells) after 1 hour that was sustained 24 hour post-treatment. Collectively, these results confirm that SB939 and 1179.4b effectively sensitise brain-metastatic mouse and human lines to radiation, resulting in induction of DNA double strand breaks and reduced ability to form colonies.

### Molecular characterisation of 4T1Br4 tumours demonstrates clinical relevance to human brain-metastatic breast cancer

We interrogated sorted tumour cells from the 4T1Br4 primary tumours versus those of highly (non-brain) metastatic 4T1 variants (4T1.2ncA5, 4T1.13ch5, 4T1BM2ch11, 4T1ch9) and an isogenic non-metastatic line (67NRch1), to identify genes associated with brain-specific metastasis. While our model clearly recapitulates the clinical brain-metastatic phenotype *in vivo*, we sought to determine whether its expression profile and underlying biology is similarly reflective of the human disease and could therefore allow identification of potential therapeutic targets. By utilising the Affymetrix Mouse Exon 1.0 ST Array with updated probeset definitions that retained only those probes mapped unambiguously to the latest genome build (Dai et al., 2005), we measured the expression of 32,722 unique ENSEMBL Gene IDs. To compare our data with human clinical cohorts and gene signatures, these IDs were converted to the HGNC Symbols of their corresponding human homologs (as per Table S1) for all subsequent analyses.

We reasoned that by identifying differentially expressed genes among brain-metastatic tumours of the 4T1Br4 model in combination with those of the Bos GSE12276 cohort, to the exclusion of genes associated with non-brain metastasis, we would derive a more brain-specific and clinically relevant gene signature with which to elucidate the underlying biology driving brain metastasis in breast cancer patients. To identify genes associated with promotion of brain-specific metastasis, we used “limma” (Ritchie et al., 2015) to measure differential expression (fold change ≥ 1.5, p ≤ 0.05) shared by tumour transcriptomes of 4T1Br4 brain metastatic versus highly metastatic (Table S2), 4T1Br4 brain-versus non-metastatic (Table S3) and Bos (GSE12276) brain-versus non-metastatic tumours (Table S4), that was not shared in 4T1 highly-versus non-metastatic tumours (Table S5). The resulting gene lists (Table S6) were intersected with VennPlex (Cai et al., 2013), yielding 30 differentially expressed gene lists among 45 distinct regions of intersection (Table S7) and revealing a 34-gene signature corresponding to the breast cancer brain-specific metastasis (BrCa BSM) gene set in region 30 (Fig. 6A*, Table S8).

**Fig. 6.**
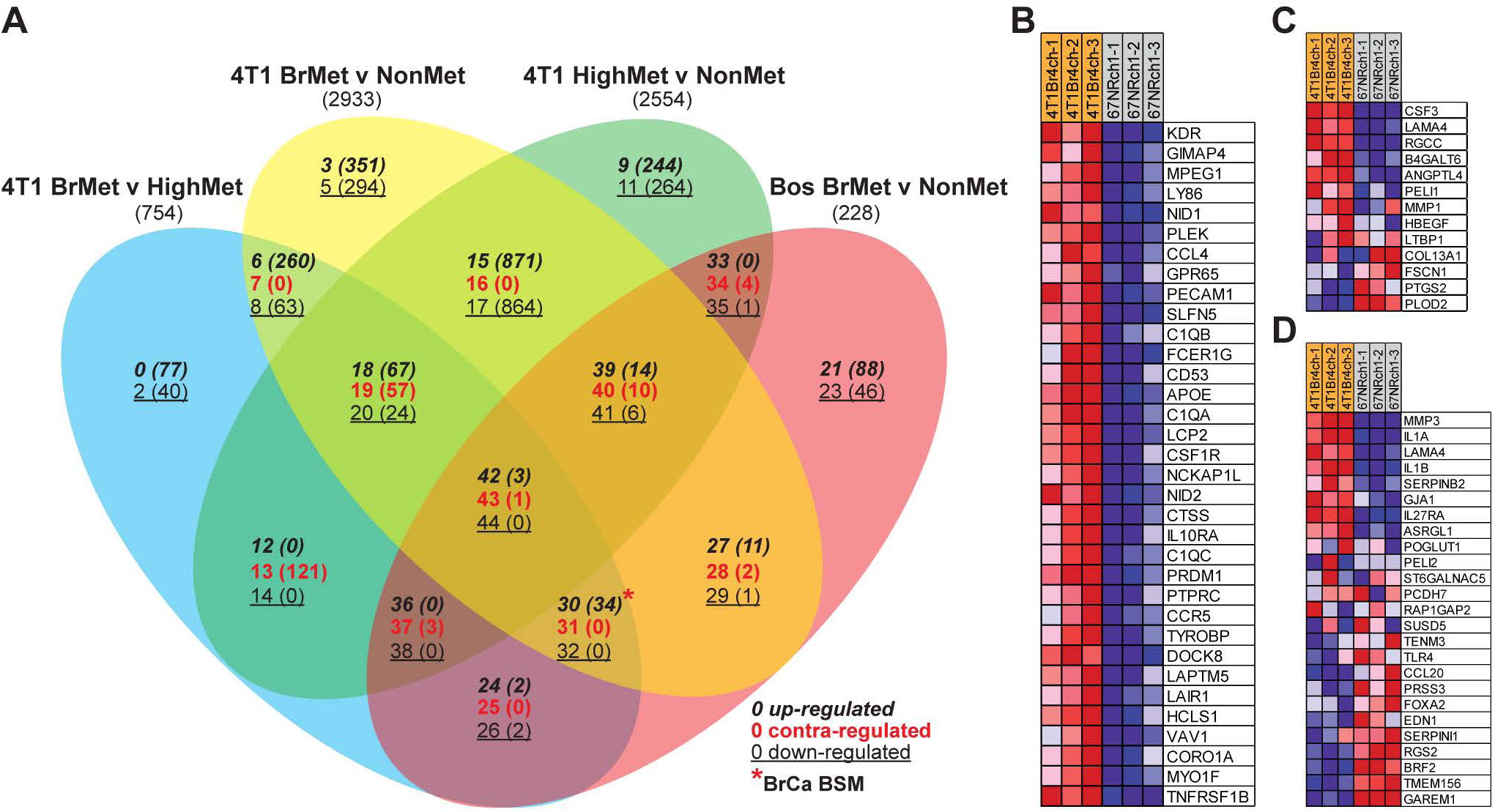
Derivation and Gene Set Enrichment Analysis of the breast cancer brain-specific metastasis gene set. (A) VennPlex was used to compare and visualise differentially expressed genes of primary breast tumours: 4T1Br4 brain-metastatic versus highly metastatic (4T1 BrMet v HighMet, blue elipse), 4T1Br4 versus non-metastatic (4T1 BrMet v NonMet, yellow elipse), 4T1 highly versus non-metastatic (4T1 HighMet v NonMet, green elipse) and GSE12276 brain-metastatic versus non-metastatic (Bos BrMet v NonMet, red elipse). Regions representing distinct gene groups are numbered, with the number of genes in that class in brackets, and indicate the order of gene sets in the extracted output. The direction of differential expression in each region is indicated, with up-regulation in *italics*, contra-regulation in red and down-regulation underlined. Region 30, corresponding to the breast cancer brain-specific metastasis (BrCa BSM) gene set is indicated by *. Heat maps generated by GSEA software indicate relative high (red) and low (blue) expression of genes in the BrCa BSM gene set (B), Bos BrMS UP gene set (C) and Bos CN34-231 BrM UP gene set (D) in 4T1Br4 brain-metastatic versus 67NR non-metastatic tumours.

To demonstrate the brain specificity of our BrCa BSM and the clinical relevance of the 4T1Br4 model, we performed gene set enrichment analysis (GSEA) (Subramanian et al., 2005) on expression datasets from the 4T1Br4 model, and human clinical cohorts GSE12276 and GSE46928 (Zhao et al., 2013) (Table S9-13), comprised of brain- and non-brain metastatic phenotypes. Specifically, we compared enrichment in brain-metastatic samples for our BrCa BSM gene set versus recently published gene sets associated with brain-metastasis derived from human breast cancer expression profiles of brain-metastatic primary tumours and cell lines ((Bos et al., 2009) Tables S4 and S7 therein). In addition, we created custom gene sets derived from differentially expressed genes of brain-versus non-brain metastatic tumours (Table S2-4, S14-18), using GSEA as a surrogate measure of the similarity and hence, the clinical relevance of the 4T1Br4 model and the BrCa BSM gene set. We found that 4T1Br4 brain-metastatic tumours were significantly positively enriched for the brain metastasis-associated gene sets from human tumours (“Bos BrMS UP”) and cell lines (“Bos CN34-231 BrM UP”) (Fig. 6B-D). Importantly, the 4T1Br4 brain-metastatic tumours were also significantly enriched for gene sets derived from up-regulated genes of all brain-metastatic samples and those of the TNBC subtype within GSE12276 (Table S19).

Next, we compared enrichment for the BrCa BSM gene set versus the other brain metastasis signatures in GSE12276 and sought confirmation that the brain-metastatic tumours of this human clinical cohort were enriched for gene sets derived from 4T1Br4 differential expression (Table S17). Significantly, enrichment for the BrCa BSM gene set in the GSE12276 brain-metastatic tumours was greater and more significant than for “Bos BrMS UP” and “Bos CN34-231 BrM UP” gene sets (NES = 3.04, FDR < 0.001, versus NES = 1.77, FDR = 0.002 and NES = 1.57, FDR = 0.010, respectively) and GSE12276 brain-metastatic tumours were indeed enriched for differentially expressed genes of 4T1Br4 versus non-metastatic tumours (Fig. S3, Table S20), consistent with the clinical relevance of 4T1Br4. To determine whether the results detailed herein were unique to 4T1Br4 and GSE12276 or may have more universal applicability, we utilised the GSE46928 cohort comprised of brain- and highly-metastatic tumours to test enrichment for corresponding gene sets (Table S18). Indeed, brain-metastatic tumours of GSE46928 were significantly enriched for several 4T1Br4 and GSE12276 differential expression gene sets and once again, BrCa BSM exhibited the greatest enrichment (Fig. S4, Table S21). Collectively, these GSEA data (Table S22) demonstrate the clinical relevance of the 4T1Br4 model on a transcriptomic level.

### Pathway analyses implicate tumour-intrinsic immune regulation and interactions with the vasculature as critical mechanisms underlying brain-specific metastasis

To gain insights into the underlying biology driving breast cancer brain-specific metastasis, we used two approaches. First, we uploaded the BrCa BSM gene set with associated fold-changes and p-values from 4T1Br4 versus non-metastatic tumours (Table S23) to MetaCore (GeneGO, Thomson Reuters) to test for enrichment in Pathway Maps, Process Networks, Diseases (by Biomarkers) and GO Processes. Overwhelmingly, immunity was the dominant theme when the uploaded dataset was intersected with curated Pathway Maps, with eight of the top ten pathways related to immune response, including the most significantly enriched pathway, “Immune response_Antigen presentation by MHC class I:cross-presentation” (Fig. 7, Table S24). Similarly, enrichment results for Process Networks (Table S25), Diseases (Table S26) and GO Processes (Table S27) were also dominated by pathways and processes involved in immunity. Clearly, tumour-intrinsic immune regulation is of paramount importance in breast cancer brain-specific metastasis.

**Fig. 7.**
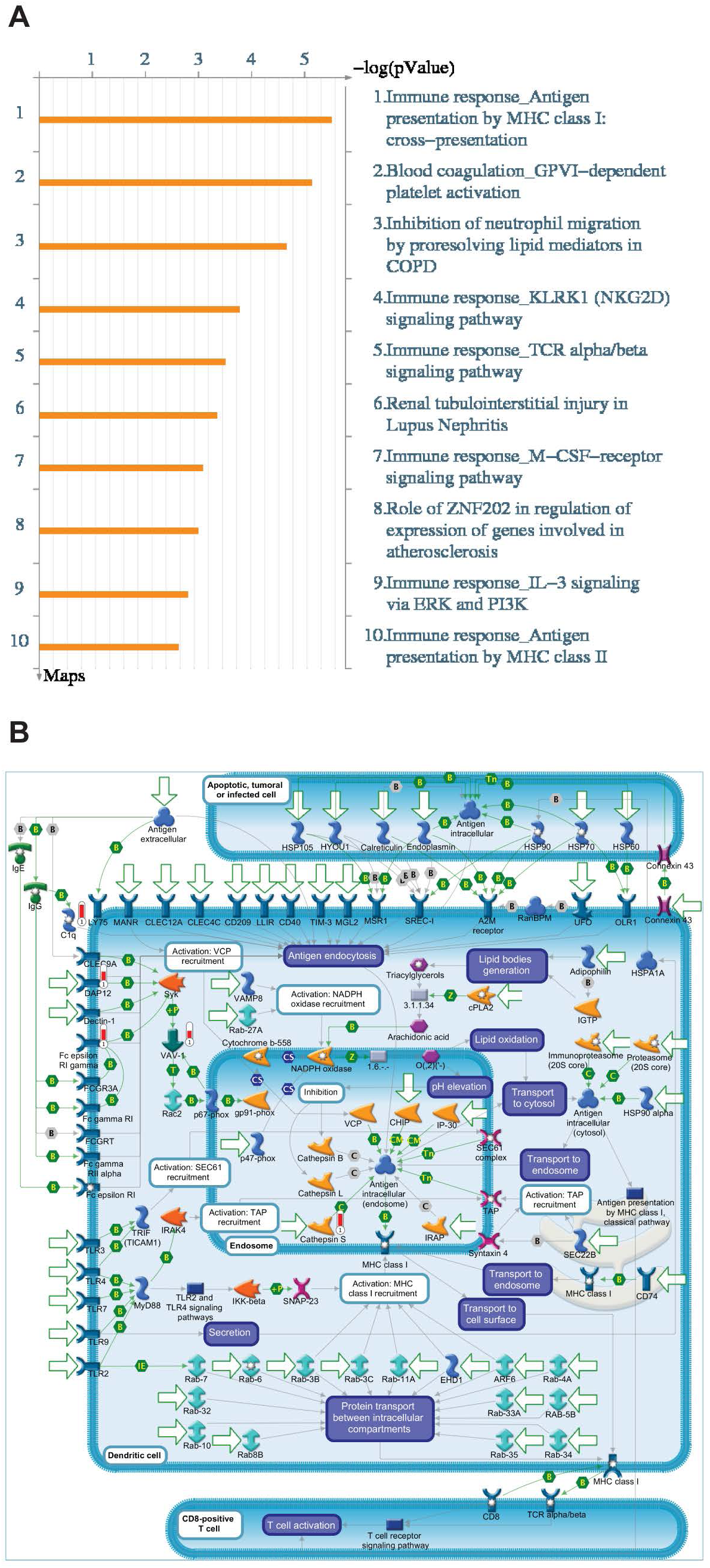
MetaCore Pathway Enrichment Analysis. The HGNC Symbols of the BrCa BSM gene set, with fold-changes and p-values from 4T1 brain-metastatic versus non-metastatic tumours, were uploaded to the MetaCore platform for Pathway Enrichment Analysis (Thomson Reuters, https://portal.genego.com/) to discover non-random functional networks associated with brain-specific metastasis. The top 10 Pathway Maps. (A) are predominantly immune-related networks, implicating tumour-intrinsic immune regulation in breast cancer brain metastasis, with “Immune response: Antigen presentation by MHC class I: cross presentation” the most significant pathway (B). The direction and degree of differential expression of genes from uploaded data is represented on the network with adjacent thermometer symbols (red, up-regulated; blue, down-regulated). For a detailed description of all MetaCore objects and symbols refer to https://portal.genego.com/help/MC_legend.pdf.

Second, we sought to identify curated Molecular Signature Database (MSigDB) gene sets (Canonical pathways, BioCarta, KEGG and Reactome; v6.1) associated with breast cancer brain-specific metastasis (BrCa BSM pathways), in an approach analogous to derivation of the BrCa BSM gene set. Specifically, GSEA was completed to detect positive enrichment of curated gene sets shared among the brain-metastatic samples of expression datasets for the 4T1Br4 model (Table S9-10) and GSE12276 (Table S11) that were not enriched in highly metastatic samples of the 4T1 (HighMet versus NonMet) expression dataset (Table S12). The curated gene sets so derived define the BrCa BSM pathways (Table 1). In support of our functional assays (Fig. 2D, E), the most significantly enriched pathway (FDR = 3.11 × 10^−3^) relates to a critical mechanism that permits tumour cell colonisation of the brain – trans-endothelial migration across the BBB (Bos et al., 2009). To prioritise potential therapeutic targets driving brain-specific metastasis, we identified 11 consensus genes contributing to enrichment in this 80-gene pathway across all brain-metastatic 4T1Br4 and GSE12276 tumours, two of which met the differential expression threshold (FC ≥ 1.5, pval ≤ 0.05) to qualify as members of the BrCa BSM gene set: *FCER1G* and *PECAM1* (Table S28). In support of the MetaCore pathway analysis, immune pathways also featured prominently among the MSigDB curated gene sets enriched in brain-metastatic tumours. Taken together, these analyses strongly implicate tumour-intrinsic immune regulation and underscore the importance of tumour cell interactions with the vasculature in breast cancer brain metastasis.

**Table 1.** GSEA enrichment report for MSigDB curated gene sets in breast cancer brain-specific metastasis expression datasets: BrCa BSM pathways.

| REACTOME, KEGG, CANONICAL PATHWAYS, BIOCARTA gene sets | SIZE | Avg ES | Avg NES | Avg FDR |
| --- | --- | --- | --- | --- |
| REACTOME_CELL_SURFACE_INTERACTIONS_AT_THE_VASCULAR_WALL | 80 | 0.61 | 2.09 | 3.11E-03 |
| KEGG_VIRAL_MYOCARDITIS | 61 | 0.61 | 2.02 | 3.29E-03 |
| REACTOME_IMMUNOREGULATORY_INTERACTIONS_BETWEEN_A_LYMPHOID_AND_A_NON_LYMPHOID_CELL | 39 | 0.70 | 2.15 | 3.74E-03 |
| KEGG_COMPLEMENT_AND_COAGULATION_CASCADES | 65 | 0.60 | 2.00 | 4.25E-03 |
| BIOCARTA_MONOCYTE_PATHWAY | 11 | 0.86 | 1.94 | 5.13E-03 |
| KEGG_INTESTINAL_IMMUNE_NETWORK_FOR_IGA_PRODUCTION | 39 | 0.68 | 2.08 | 6.34E-03 |
| REACTOME_IMMUNE_SYSTEM | 763 | 0.44 | 1.94 | 8.11E-03 |
| KEGG_NATURAL_KILLER_CELL_MEDIATED_CYTOTOXICITY | 95 | 0.54 | 1.94 | 1.10E-02 |
| PID_GMCSF_PATHWAY | 35 | 0.64 | 1.91 | 1.13E-02 |
| KEGG_CELL_ADHESION_MOLECULES_CAMS | 113 | 0.55 | 2.04 | 1.26E-02 |
| REACTOME_ANTIGEN_PROCESSING_CROSS_PRESENTATION | 66 | 0.55 | 1.85 | 1.92E-02 |
| KEGG_PRION_DISEASES | 33 | 0.61 | 1.80 | 2.16E-02 |
| BIOCARTA_IL10_PATHWAY | 17 | 0.75 | 1.88 | 2.23E-02 |
| REACTOME_RESPONSE_TO_ELEVATED_PLATELET_CYTOSOLIC_CA2 | 72 | 0.53 | 1.80 | 2.40E-02 |
| PID_IL4_2PATHWAY | 60 | 0.55 | 1.80 | 2.50E-02 |
| KEGG_ECM_RECEPTOR_INTERACTION | 83 | 0.51 | 1.78 | 2.58E-02 |
| REACTOME_SIGNALING_BY_ILS | 100 | 0.50 | 1.80 | 2.66E-02 |
| REACTOME_PLATELET_ACTIVATION_SIGNALING_AND_AGGREGATION | 187 | 0.47 | 1.82 | 2.68E-02 |

## DISCUSSION

We have described the successful development of a new model of breast cancer brain metastasis that incorporates several elements of clinical relevance including robust spontaneous spread from the mammary gland to the brain in immune competent animals. TNBC is characterised by earlier onset of metastasis compared to other molecular subtypes and high incidence of brain metastasis, often with extensive extracranial involvement as well. While 4T1Br4 and parental 4T1 tumours have a pattern of metastatic spread that is similar to that reported for human TNBC (Liedtke et al., 2008), we found that 4T1Br4 tumours are selectively more metastatic to the brain compared to parental 4T1 tumours. To the best of our knowledge, 4T1Br4 tumours show the highest incidence of spontaneous breast cancer brain metastases among all syngeneic mouse models described so far. While other investigators have reported on the use of brain-tropic 4T1 cell variants with distinct growth and phenotypic characteristics (Erin et al., 2013; Lockman et al., 2010), it is not clear from these studies whether the incidence of spontaneous brain metastasis in these models is sufficiently robust for preclinical evaluation of new therapies. Optimisation of the 4T1Br4 model here required *in vivo* serial enrichment, clonal selection of metastatic variants, resection of the primary tumour and injection of fewer cells into the mammary fat pad. These modifications had a cumulative effect on the incidence of brain metastasis and prevented early termination of experiments due to unethical primary tumour size and excessive lung metastatic burden as noted by others (Bailey-Downs et al., 2014; Zhang et al., 2015).

Clonal selection also identified significant heterogeneity in brain metastatic potential between individual 4T1Br4 clones and the bulk 4T1 parental cell population. The concept of tumour heterogeneity is now well recognised and the genomic heterogeneity between matched primary tumours and brain metastases suggests that brain metastases are likely to arise from a minority of cells present in the primary tumour (Ding et al., 2010). Molecular and functional heterogeneity has been demonstrated also in the 4T1 model (Martelotto et al., 2014) and was exploited by our group previously to derive highly bone-metastatic (but not brain-metastatic) 4T1 variants (Kusuma et al., 2012). It is also noteworthy that differences in the anatomical location and number of brain lesions were observed between spontaneous and experimental metastasis assays. While experimental metastasis assays gave rise to multiple metastatic foci scattered through the cerebrum, cerebellum, leptomeninges and olfactory lobes, only one or two brain lesions most commonly formed in the cerebrum in the spontaneous metastasis assay. The latter appear to be more representative of the pattern of brain metastasis observed in the clinic (Delattre et al., 1988). These differences can be explained in part by the large number of cells injected directly into the vasculature in experimental metastasis assays, resulting in multiple clones successfully homing to and colonising the brain, including those that may not possess all functional attributes required for spontaneous brain metastasis. Collectively, these observations demonstrate the greater clinical relevance of the spontaneous metastasis assay.

Functionally, 4T1Br4 cells demonstrated enhanced migratory activity and ability to adhere to and migrate through brain microvascular endothelial cells. These properties are likely to contribute to the brain selectivity of 4T1Br4 cells *in vivo* and are consistent with observations in melanoma and other breast cancer brain metastasis models (Bos et al., 2009; Cruz-Munoz et al., 2008). Three genes, Cox2, HBEGF and ST6GALNAC5, identified by Bos and colleagues (Bos et al., 2009) were shown to promote trans-endothelial migration in breast cancer cell lines. While our study did not specifically investigate the role of these genes, our analyses implicated several genes associated with endothelial cell adhesion in brain-specific metastasis, including *PECAM1*, a transmembrane protein directly linked to trans-endothelial migration (Muller et al., 1993) across the BBB (Akers et al., 2010; Winger et al., 2014). PECAM1 is expressed on several hemopoietic lineages and is one of the quintessential markers of the vasculature in immunohistology (Watt et al., 1995). While we cannot completely rule out stromal and tumour-infiltrating cells as contaminants, our expression profiling utilised tumour cells FACS-isolated by their endogenous mCherry fluorescence. Hence, any host-derived co-purifying cells would be rare and unlikely to provide a significant signal. Indeed, PECAM1 has been detected on several rodent and human solid tumour cell lines, including breast carcinoma MCF7 (Maniotis et al., 1999; Tang et al., 1993), and human brain glioblastoma (Aroca et al., 1999).

In addition, our analyses revealed the importance of tumour-intrinsic immune regulation in brain-specific metastasis, due in no small part to the many differentially expressed genes traditionally viewed as leukocytic markers, *FCER1G* chief among them. These observations are not without precedent, as several studies have reported similar “inappropriate gene expression” of leukocytic markers in metastatic breast and other cancers (Tarin, 2012). Hence, tumour-specific FCER1G and PECAM1 may represent *bona fide* therapeutic targets that warrant further evaluation.

The high incidence of spontaneous 4T1Br4 brain metastasis makes this model ideally suited for pre-clinical evaluation of new therapeutics. Our *in vitro* investigation showed that SB939 and 1179.4b are significantly more potent than SAHA at inhibiting the proliferation/survival of 4T1Br4 and human MDA-MB-231Br brain-tropic lines, in agreement with the higher potency of these compounds in other tumour lines (Kahnberg et al., 2006; Novotny-Diermayr et al., 2010). Whilst we have yet to investigate the specific mechanisms and signalling pathways by which SB939 and 1179.4b exert their inhibitory effects, our data indicate that the significant reduction in tumour growth may be mediated in part through inhibition of histone H3 deacetylation as observed *in vitro* and *in vivo* and/or by induction of DNA damage as evidenced by the prolonged (≥ 24 hours) increase in the number of γH2AX foci seen in MDA-MB-231Br cells.

Our study is the first to show that SB939 and 1179.4b significantly inhibit the orthotopic growth of brain-metastatic breast tumours. Interestingly, acetylation of histone H3 after single treatment of 4T1Br4 cells with SB939 or 1179.4b *in vitro* was reversible, with rapid deacetylation upon removal of SB939 (<1 hour) or 1179.4b (<8 hours). This contrasts with the high acetylation level observed in primary tumours analysed after the last treatment. These observations indicate that SB939 and 1179.4b most likely accumulate in 4T1Br4 tumours and induce sustained acetylation of histone H3 following repeated dosage, as reported for SB939 in pre-clinical colorectal cancer models and in patients with solid tumours (Novotny-Diermayr et al., 2010; Yong et al., 2011). The effect of 1179.4b on histone H3 acetylation has not been reported previously. Additional studies are warranted to evaluate the complete pharmacokinetic/pharmacodynamic properties of 1179.4b *in vivo*.

SB939 and 1179.4b also inhibited spontaneous bone metastasis in these experiments. Given that SB939 and 1179.4b were administered in a neo-adjuvant setting without tumour resection and inhibited primary tumour growth, it is unclear whether inhibition of 4T1Br4 bone metastasis is attributable to lower metastatic dissemination from smaller tumours or a direct effect of these inhibitors on the growth of metastases in bone or both. Interestingly, Novotny-Deirmayr and colleagues (Novotny-Diermayr et al., 2011) noted that acetylation of histone H3 in bone marrow becomes detectable after prolonged treatment with SB939 in a model of acute myeloid leukaemia. Thus, it is reasonable to suggest that SB939 may also accumulate in 4T1Br4 bone metastases and inhibit their growth directly. Moreover, recent studies showed that 1179.4b potently inhibits osteoclast bone resorption activity *in vitro* and suppresses bone loss in an experimental model of periodontitis (Cantley et al., 2012). Osteoclasts are important mediators of osteolytic activity and bone loss associated with the development of breast cancer bone metastases. Together, these studies and ours suggest that, in addition to their direct effect on tumour cells, inhibition of osteoclast function by pan-HDACi such as 1179.4b may further contribute to inhibiting osteolytic bone metastases. Future studies will be required to confirm this possibility.

In the adjuvant setting, where therapy was initiated after removal of the primary tumour, SB939 and 1179.4b dramatically reduced the number of detectable brain lesions. It was not possible to determine whether spontaneous brain micrometastases were already present prior to the beginning of HDACi treatment due to the limited mCherry fluorescence signal in micro-lesions. Thus, it is unclear whether these HDACi directly inhibited established brain metastases or effectively killed circulating breast cancer cells in the vasculature before they home to and colonise the brain (preventive effect). Structurally, SB939 meets several requirements for BBB penetration including low molecular weight (≤ 500 Da) and oil/water distribution coefficient (LogP ≤ 5) and absence of efflux ratio indicating that it is unlikely to be a P-glycoprotein transporter substrate (Jayaraman et al., 2011). However, the lack of effect of SB939 on experimental brain metastasis indicates that it may not reach sufficient concentrations in the brain (or other visceral metastases) to achieve clinical significance. Whilst 1179.4b partially inhibited experimental brain metastasis, its predicted limited permeability across the BBB suggests that its effect is likely to be primarily preventive. In support of this, we found that the number of viable CTCs in blood as well as ovary, kidney and adrenal metastases were reduced in 1179.4b-treated mice. However, given the high potency of 1179.4b compared to SAHA or SB939, we cannot completely rule out that low levels of this inhibitor may be sufficient to exert a direct effect on brain metastases in which the BBB is disrupted, as seen in some TNBC patients (Yonemori et al., 2010).

Our preliminary evaluation of SB939 and 1179.4b radio-sensitising properties indicates that both inhibitors sensitise 4T1Br4 and MDA-MB-231Br cells to radiation *in vitro*. To our knowledge, this is the first report demonstrating the radio-sensitising properties of SB939 and 1179.4b in brain-metastatic TNBC models. SB939 or 1179.4b alone increased the expression of γ-H2AX at both 1 and 24 hour time points. Sustained induction of γ-H2AX by the combination of either SB939 or 1179.4b and low dose radiation (1 Gy) indicates that extensive DNA damage occurs under those conditions and contributes to the anti-tumour effect of SB939 and 1179.4b. These findings provide a strong rationale for further studies exploring the efficacy of therapies combining WBRT with either of these or other HDACi for the treatment of brain metastasis.

### Conclusions

The 4T1Br4 model of brain-metastatic breast cancer closely recapitulates the human disease at phenotypic, functional and transcriptomic levels. Therefore, we believe that our model is clinically relevant and an ideal system in which to identify potential therapeutic targets and evaluate the efficacy of pharmacological and/or genetic interventions. Notably, expression profiling and related analyses demonstrated the critical role of tumour-intrinsic immune regulation in brain-specific metastasis, underscoring the importance of using immune-competent models as tools for discovery and for translational studies that may incorporate immunotherapeutic strategies. These data also implicated tumour cell surface interactions with the vasculature in successful colonisation of the brain, in support of previous studies and functional assays herein. Significantly, curated gene set enrichment analysis identified several potential therapeutic targets for future evaluation, with FCER1G and PECAM1 emerging as promising candidates. Evaluation of HDACi efficacy in this model demonstrated partial efficacy of SB939 and the more potent 1179.4b against mammary tumours and metastases, including to the brain. However, these compounds are unlikely to be sufficient alone to fully control advanced metastatic disease and will require combination therapies to achieve clinically relevant disease control, in particular brain metastasis. Several clinical trials combining HDAC inhibitors and radiation therapy are now underway or in recruitment phase, some aimed specifically at treating brain metastases from solid tumours (Groselj et al., 2013). Our preliminary assessment of the radio-sensitising properties of SB939 and 1179.4b *in vitro* warrants further investigation of their therapeutic potential in combination with WBRT against brain metastasis.

## MATERIALS AND METHODS

### Cell culture and reagents

The 4T1 and 67NR mouse mammary carcinoma cell lines were obtained from Dr. F. Miller (Karmanos Cancer Institute, Detroit, MI, USA) and cultured as described previously (Kusuma et al., 2012). Highly metastatic (4T1.13ch5, 4T1BM2ch11, 4T1.2ncA5, 4T1ch9) and non-metastatic clones with stable expression of mCherry were derived as described (Carter et al., 2015; Chang et al., 2015; Lelekakis et al., 1999; Martin et al., 2017). MDA-MB-231Br cells were kindly provided by Prof J. Massague (Memorial Sloan Kettering Cancer Center, NY, USA) and bEnd.3 murine brain endothelial cells were a generous gift from Dr. R. Hallmann (Münster University, Münster, Germany). These lines were maintained in Dulbecco’s Modified Eagle’s Medium (DMEM) supplemented with 10% foetal bovine serum (FBS), 2 mM L-glutamine, 1 mM sodium pyruvate and 1% penicillin-streptomycin at 37°C with 5% CO_2_. The HCACi SB939 (Procinostat in Wang et al., 2011) and 1179.4b (compound 52 in Kahnberg et al., 2006) were synthesised and purified as described. Stocks solutions were prepared in DMSO and diluted as appropriate for *in vitro* and *in vivo* experiments. For animal studies, SB939 and 1179.4b were prepared in saline, 30% polyethylene glycol, 10% v/v dimethyl sulfoxide and administered intraperitoneally.

### Animal studies

All procedures involving mice conformed to National Health and Medical Research Council animal ethics guidelines and were approved by the Peter MacCallum Animal Ethics & Experimentation Committee (Ethics E507 and E509).

### Tumour growth and metastasis assays

Tumour cells (2 × 10^4^ in 20 µL of phosphate-buffered saline (PBS)) were injected into the 4th mammary fat pad of 6 to 8-week-old female BALB/C mice and tumour growth monitored thrice weekly with electronic callipers. Tumour growth/volume was calculated using the formula (length X width^2^)/2. Mice were euthanised by overdose of isoflurane inhalation after approximately 30 days. Where indicated, primary tumours were resected when they reached 0.4 - 0.5g and the mice euthanised as above ~3 weeks post-tumour resection or earlier if mice showed signs of distress due to metastatic disease. Lungs, spines, femurs and brains, were harvested and processed for histological examination and immunohistochemical staining or for real-time genomic qPCR analysis of relative tumour burden (RTB) using Taqman chemistry (PE Biosystems, Foster City, CA, USA) as described previously (Carter et al., 2015; Denoyer et al., 2014). For experimental metastasis assays, cells were inoculated directly into the left cardiac ventricle as described previously (Martin et al., 2017). Metastasis was scored by IHC staining of cytokeratin^+ve^ cells (3 step sections/brain, 200 µm apart) and quantitated using Aperio ImageScope version 12.3.2.8013.

When comparing metastatic burdens from two cancer cell lines, the RTB values were adjusted to relative gene copy number calculated from the ratio of signals obtained from genomic DNA purified from each cell line in culture. Primers and probes were designed using Primer Express (Applies Biosystems, Foster City, CA, USA): mCherry (forward, 5’-GACCACCTACAAGGCCAAGAAG-3’; reverse, 5’-AGGTGATGTCCAACTTGATGTTGA-3’; probe, 6FAM-CAGCTGCCCGGCGCCTACA-TAMRA), vimentin (forward, 5’-AGCTGCTAACTACCAGGACACTATTG-3’; reverse, 5’-CGAAGGTGACGAGCCATCTC-3’; probe, VIC-CCTTCATGTTTTGGATCTCATCCTGCAGG-TAMRA). Reactions were performed on an ABI Prism 7000 thermocycler. RTB in an organ was calculated using the formula: RTB = 10000 /(2^ΔCT^), where ΔCT = CT (mCherry) – CT (vimentin). Where indicated, the presence of brain and lung macro-metastases was detected by fluorescence imaging of mCherry marker using a Maestro™ In-Vivo Imaging System (CRi).

### Immunohistochemical (IHC) staining

Tissues were dissected, fixed in 10% buffered formalin and processed for paraffin embedding. Serial sections from primary tumour (4 µm) or brain (6 µm) were stained with haematoxylin and eosin (H&E) for morphology or subjected to standard IHC staining (Martin et al., 2017). Briefly, paraffin sections were de-waxed in xylene and rehydrated in graded ethanol solutions prior to heat induced epitope retrieval in 10mM citrate buffer (pH 6.0). The following primary antibodies were used: ERα (DAKO, cat. #M7047), PR (Santa Cruz Biotech, cat. # SC-538), HER2 (Calbiochem, cat. #OP15L), pan cytokeratin (SIGMA, cat #C1801), Ki-67 (Abcam, cat. #Ab15580) and anti-GFAP (DAKO, cat #Z0334). Specific binding was detected using appropriate biotin-conjugated secondary antibodies and ABC reagents (Vectastain kit #PK6100, Vector Laboratories). Staining was visualised with 3,3’-diaminobenzidine (DAB) and a haematoxylin counterstain.

### SDS-PAGE and acetylated histone H3 immunoblotting

Whole cell lysates from control and HDACi-treated sub-confluent cells cultures or crushed frozen tumours were prepared in ice-cold RIPA buffer [10 mM Tris-HCl (pH 7.4), 5 mM EDTA (pH 8), 1% (v/v) Nonidet P-40, 0.5% (v/v) sodium deoxycholate, 0.1% (v/v) SDS] supplemented with protease inhibitor cocktail (ROCHE, cat. #04693132001). Lysates were sonicated four times for 30 sec, cleared by centrifugation at 13,000 × g for 15 min at 4°C and the supernatants stored at -20°C.

Western blotting was performed using standard methodology as described previously (Denoyer et al., 2014). Briefly, 40 µg of proteins were separated on a 15% SDS-PAGE gel and transferred to a PVDF membrane. The membrane was blocked with 10% (w/v) non-fat dried milk in PBS containing 0.05% Tween-20 for 1 hour at room temperature and incubated with anti-acetylated histone H3 antibody (Millipore, cat. #06-599, 1:10,000 dilution) overnight at 4°C. After 3 washes with wash buffer (0.025% Tween-20 and 0.1% BSA in PBS) for 10 min, the membrane was incubated for 1 hour with an appropriate horseradish peroxidase-conjugated secondary antibody in wash buffer. Specific protein bands were detected using enhanced ECL reagents and Super RX film or ChemiDoc™ MP System (Biorad). An anti-GAPDH antibody (Abcam, cat. #Ab8245, 1/10000 dilution) was used as loading control.

### Preparation of brain conditioned medium (BCM)

Newborn pups from BALB/C mice were rinsed quickly with 70% alcohol and whole brains removed under sterile conditions. Brains were rinsed twice quickly with 20 mL PBS containing 2% penicillin-streptomycin and fluconazole (6 µg/mL), minced with a scalpel blade and cultured in DMEM supplemented with 20% FBS, 1mM sodium pyruvate, fluconazole (6 mg/L) and 2% penicillin-streptomycin at 37°C with 5% CO_2_. The medium was changed every 2-3 days until subconfluent. Adherent cells were serum-starved overnight in 15 mL α-MEM serum free medium (SFM) supplemented with 0.05% bovine serum albumin (BSA), 2 mM L-glutamine, 1 mM sodium pyruvate and 1% penicillin-streptomycin and the medium replaced with 10 mL of fresh SFM. Serum-free BCM was collected after 48 hours, centrifuged at 1200 rpm, at 4°C, for 5 min and stored at -80°C until used. BCM was stable for at least 3 months at -80°C.

### Colony formation assay

Colony formation assays were done as described previously with minor modifications (Martin et al., 2017). Briefly, 4T1Br4 cells (100/well) were seeded in triplicate wells of a 6-well plate in SFM supplemented or not with 1% FBS, 50% BCM or both. Cells were fixed in methanol, stained with crystal violet and the number of colonies (>50 cells) formed after 10 days counted manually. The effect of histone deacetylase inhibitors on cell survival was determined in the same assay in 4 mL of growth medium containing 0.1% DMSO (vehicle), 1 μM SAHA, 1 μM SB939 or 1 μM 1179.4b for 10 days at 37°C. For quantitation of viable CTCs from control and HDACi-treated mice, blood was collected by cardiac puncture, cleared of erythrocytes by hypotonic shock and resuspended in complete medium. Thirty µL, 100 µL or 300 µL of cell suspension were seeded in triplicate in 6 well plates containing 4 mL of growth medium and incubated at 37°C. Colonies (> 50 cells) formed after 10 days were counted and the data expressed as mean number of viable CTCs/mL of blood. Statistical difference between groups was analysed using a Mann-Whitney nonparametric test when comparing two groups or a one-way ANOVA, Bonferroni post-test when comparing multiple groups; *p* < 0.05 was considered significant.

### Proliferation assay and determination of inhibitory concentration (IC_50_)

Cell proliferation was measured using a sulforhodamine B (SRB) colorimetric assay as described previously (Denoyer et al., 2014). Proliferation was measured over 5 days in SFM supplemented with 50% v/v BCM or 1% FBS or both. Initial cell density was 5 x 10^2^ cell/well in 96 well plates (6 replicate wells per condition). Assays were completed three times and the data presented as the mean ± SD of 6 replicates/condition from a representative experiment. Statistical significance was determined using a two-way ANOVA; p < 0.05 was considered statistically significant. IC_50_ values were determined in the same assay over 3 days with an initial cell density of 1 × 10^3^ cells/200 µL/well as described (Martin et al., 2017). IC_50_ values were calculated using Hill’s equation in the GraphPad Prism 6.0 software.

### Migration and invasion assays

Chemotactic migration assays were completed in triplicate Transwell polycarbonate membrane inserts (8 µm pore size) (Corning Inc., Life Sciences, NY, USA) as described previously (Sloan et al., 2006). Tumour cell migration to the underside of porous membranes was measured after 4 hours of incubation at 37°C. Cells on the underside of membrane were fixed in 10% neutral buffered formalin, permeabilised for 5 min in 0.1% Triton X-100 and stained with 0.5µg/mL 4′,6-diamidino-2-phenylindole (DAPI). For invasion assays, tumour cells (1x 10^5^) were embedded in 100 µL of a 1:1 mixture of SFM and Matrigel (BD Biosciences, CA, USA) in the upper wells and allowed to invade and migrate towards SFM (control) or 50% BCM. The number of cells on the underside of the porous membrane was scored after 18 hours of incubation at 37°C, as described for migration assays. All migration and invasion experiments were repeated three times. Three random fields per membrane were photographed on an Olympus BX61 microscope with 20 × magnification and the number of migrated/invaded cells counted using Metamorph Image Analysis software (Molecular Devices, CA, USA). The results are expressed as the mean number of migrated/invaded cells per field ± SD of nine replicates (3 replicate wells × 3 field of view per condition) from a representative experiment (n = 3). The statistical differences were analysed using a Mann-Whitney nonparametric test when comparing two groups or a one-way ANOVA, Bonferroni post-test when comparing multiple groups; p < 0.05 was considered significant.

### Adhesion to endothelial cell and trans-endothelial cell migration assays

Adhesion assays were performed in 96-well culture plates as described previously (Sloan et al., 2006). In brief, bEnd.3 cells (1 × 10^5^/200 µL) were seeded in triplicate wells of 96-well plates and incubated at 37°C to allow formation of a monolayer. Monolayers were washed twice with PBS before addition of calcein-labelled tumour cells (4 × 10^4^) in 100µL SFM (Calcein AM, ENZO Life Sciences Inc., NY, USA). The plates were incubated for 10 min at 4°C and further incubated for 30 min at 37°C. Non-adherent cells were removed by gentle washing twice with PBS and once with Tris-buffered saline (pH 7.4) supplemented with 2 mmol/L CaCl_2_ and 1 mmol/L MgCl_2_. Fluorescence was measured using a Bio-Rad Molecular Imager FX reader and specific adhesion calculated from a standard curve made up of 0, 25, 50, 75 and 100 µL of calcein-labelled cell lysates derived from the initial cell suspension. For trans-endothelial cell migration assays, bEnd.3 cells (1x 10^5^) were seeded in triplicate Transwell inserts as above in complete DMEM medium supplemented with 10% FBS for 24 hours. Monolayers were washed as above and calcein-labelled tumour cells (2 × 10^5^ cells/200 µL SFM) were added to the upper well. Six hundred µL of SFM alone or SFM supplemented with 5% FBS and laminin-511 (1 µg/mL) were added to the lower chamber. After 48 hours incubation, migrated tumour cells (calcein AM positive) on the underside of the porous membrane were processed for microscopy as described for the migration assay and counted. All adhesion to endothelial cell and trans-endothelial cell migration experiments were repeated three times and the results are presented as the mean percentage of total cell ± SD from a representative experiment. The statistical differences were analysed using a Mann-Whitney nonparametric test when comparing two groups or a one-way ANOVA, Bonferroni post-test when comparing multiple groups; p < 0.05 was considered significant.

### Radiosensitisation and **γ**-H2AX immunofluorescence staining

4T1Br4 cells (100/well) were seeded in complete medium in 6 cm culture dishes and allowed to adhere for 6 hours at 37°C. Adherent cells were treated with SB939 (530 nM) or 1179.4b (70 nM) for 24 hour followed by irradiation at 2, 4 or 6 Gy. Colonies were quantitated after 10 days at 37°C as described above and the results expressed as mean surviving fraction ± SD of triplicate wells from a representative experiment (n = 3). p-values were calculated using a one-way ANOVA with a Bonferroni post-test; p < 0.05 was considered significant.

The radiosensitising effect of SB939 or 1179.4b on MDA-MB-231Br cells was determined by detection of γ-H2AX. Cells (4 × 10^4^ cells/500 μl/well) were seeded in chamber slides in DMEM medium supplemented with 10% FBS and 1% penicillin-streptomycin and incubated for 18 hours at 37°C. The medium was removed and replaced with 500 μl of growth medium containing either 360 nM SB939 or 50 nM 1179.4b, corresponding to their respective IC_50_. Following 24 hours incubation at 37°C, cells were irradiated with 1 Gy. One or 24 hours after irradiation, cells were rinsed in PBS and fixed with 4% PFA for 30 minutes, placed in 70% ice-cold ethanol and stored at 4°C. For γ-H2AX staining, cells were washed with PBS for 15 min at room temperature and blocked with 8% BSA in PBS containing 0.5% Tween-20 and 0.1% Triton X-100 (PBS-TT) for 30 min at RT. Slides were incubated with an anti-γ-H2AX antibody (Abcam, cat. #Ab11174, 1:500 dilution in 1% BSA in PBS-TT) for 2 hours at room temperature under a humidified atmosphere before washing three times with PBS for 5 minutes and incubated with a secondary antibody (Alexa fluor 488 conjugated goat anti-rabbit IgG, 1:500 dilution in 1% BSA in PBS-TT) for 1 hour at room temperature. Slides were washed three times as above and mounted with a Vectashield mounting medium containing DAPI. The results are expressed as the mean number of cells positive for γ-H2AX/per field ± SD of nine fields of view (3 replicates × 3 fields of view per condition) from a representative experiment (n = 3). Statistical differences were analysed using a one-way ANOVA, Bonferroni post-test; p < 0.05 was considered significant.

### Tumour processing, RNA isolation and expression profiling

Tumours (~0.5g) were resected, minced and digested in 10 mL DMEM with 750U/mL collagenase Type I (Worthington) for 45 min at 37°C with 180 rpm shaking, whereupon 10 mL ice-cold DMEM with 20% FBS were added to stop digestion. Following debris removal through 100 µm and 40 µm strainers and red blood cell lysis (75 mM NH4Cl, 1 mM KHCO3, 100 nM EDTA, 5 min at 23°C), viable (30 nM Sytox Green-negative) mCherry^+ve^ tumour cells were isolated by FACS and cell pellets snap-frozen in liquid nitrogen. Total RNA was isolated with the mirVana miRNA isolation kit (Ambion), followed by rRNA depletion with RiboMinus (Invitrogen), DNA removal with Turbo DNA-free (Ambion, rigorous protocol) and concentration with the RNeasy MinElute kit (QIAGEN). RNA integrity was assessed with the Agilent 2100 Bioanalyzer and samples were hybridised to Affymetrix Mouse All Exon 1.0 ST microarrays, washed, stained and scanned according to manufacturer’s protocols. The raw microarray signal (CEL) and associated files have been deposited in NCBI’s Gene Expression Omnibus (Edgar et al., 2002) and are accessible through GEO Series accession number GSE111489 (https://www.ncbi.nlm.nih.gov/geo/query/acc.cgi?acc=GSE111489).

CEL files and associated clinical data for GSE12276 (Bos et al., 2009) and GSE46928 (Zhao et al., 2013) were downloaded from the ArrayExpress Archive of Functional Genomics Data (http://www.ebi.ac.uk/arrayexpress) (Kolesnikov et al., 2015). Microarrays were analysed with the R statistical computing language (https://www.R-project.org/) and all R software packages were sourced from the Bioconductor repository (Gentleman et al., 2004). CEL files were processed with the “affy” R package (Gautier et al., 2004) using the custom chip definition files (CDFs) “hgu133plus2hsensg” and “hgu133ahsensg”, for GSE12276 and GSE46928, respectively; “moex10stmmensg” CDF was utilised for mouse exon arrays (V22, Dec 2017) (Callagy et al., 2005). Outlier arrays (GSE12276: GSM308285, GSM308293, GSM308316, GSM308339, GSM308340, GSM308358, GSM308362, GSM308364, GSM308371, GSM308376, GSM308406, GSM308420, GSM308424, GSM308429, GSM308444, GSM308453, GSM308457; GSE46928: none) were identified with the “arrayQualityMetrics” R package (Kauffmann et al., 2009) and removed from subsequent analyses. Samples were normalised with justRMA and median polish probe summarization (Irizarry et al., 2003) and differentially expressed genes (fold change ≥ 1.5, p ≤ 0.05) were identified using “limma” (Ritchie et al., 2015) to compare expression profiles of tumours grouped according to metastatic potential (Table S2-S5). Human genomic data was sourced from Ensembl using the online BioMart data mining tool (Smedley et al., 2009); mouse genomic data and human-murine orthologs (Table S1) were sourced from the Mouse Genome Informatics (MGI) repository (http://www.informatics.jax.org/). Expression datasets were joined to corresponding genomic datasets on common fields using the Galaxy (Giardine et al., 2005) platform.

### Derivation of the breast cancer brain-specific metastasis (BrCa BSM) gene set

The intersection of differentially expressed gene lists from brain- (“BrMet”) versus highly- (“HighMet”) and non-metastatic (“NonMet”) tumours was implemented with VennPlex software (Cai et al., 2013) to identify consensus ‘driver’ genes associated with brain metastasis from the 4T1Br4 and Bos GSE12276 cohorts. Specifically, the genes of interest corresponded to those common to 4T1 brain-metastatic versus highly-metastatic (“BrMet v HighMet”, Table S2), 4T1 brain-versus non-metastatic (“BrMet v NonMet”, Table S3), Bos brain-metastatic versus local/non-recurring (“Bos BrMet v NonMet”, Table S4) and not including 4T1 highly-versus non-metastatic (“HighMet v NonMet”, Table S5) differentially expressed genes. The intersection of these gene lists (Table S6) resulted in 45 distinct regions (Table S7), with “region_30” corresponding to the BrCa BSM gene set (Fig. 6A, “*”; Table S8).

### Gene set enrichment analysis and derivation of BrCa BSM pathways & targets

Gene set enrichment analysis (GSEA) (Subramanian et al., 2005) was utilised to enable detection of co-ordinated expression across sets of genes in brain-versus non-brain metastatic tumour expression datasets (log2 RMA-normalised expression values, Table S9-13). Phenotype labels were associated with each sample according to our site-specific metastasis data or from corresponding clinical data available from ArrayExpress. The GSEA desktop application and curated gene sets were downloaded from the Broad Institute GSEA site (http://software.broadinstitute.org/gsea/index.jsp). Analyses were performed with 1,000 gene set permutations using native HGNC gene symbol identifiers. Custom gene sets for brain-versus non-brain metastatic samples of the 4T1 and Bos cohorts (Table S16-18) were constructed from corresponding lists of differentially expressed genes (Table S2-4, S14-15); TNBC samples within GSE12276 were identified as per associated hormone receptor clinical data. The “Bos BrMS UP” and “Bos CN34-231 BrM UP” brain metastasis gene sets were derived from (Bos et al., 2009), Supplementary Tables 4 and 7, respectively. Gene set enrichment reports for 4T1 (Table S19), GSE12276 (Table S20) and GSE46928 (Table S21) expression datasets correspond to positive enrichment of the given gene sets in the brain-versus non-brain metastatic samples, ordered by FDR q-val, then normalized enrichment score (NES); GSEA analyses for all cohorts were summarized (Table S22). To identify positive enrichment for curated MSigDB gene sets (Canonical pathways, BioCarta, KEGG and Reactome; MSigDB v6.1) associated with breast cancer brain-specific metastasis (BrCa BSM pathways), GSEA was used to identify curated gene sets enriched in the brain-metastatic samples of expression datasets for 4T1 BrMet versus HighMet (Table S9), 4T1 BrMet versus NonMet (Table S10) and Bos BrMet versus NonMet (Table S11) that were not enriched in highly metastatic samples of the 4T1 HighMet versus NonMet expression dataset (Table S12). The unique curated gene sets commonly enriched in brain-metastatic samples across 4T1 and GSE12276 cohorts correspond to the BrCa BSM pathways (Table 1).

### GeneGO MetaCore pathway analysis

To gain insights into the functional pathways and processes underlying the BrCa BSM gene set, the HGNC Symbols, fold-changes and p-values of the 34 up-regulated genes in 4T1Br4 brain-versus non-metastatic tumours (Table S23) were uploaded to MetaCore (Thomson Reuters, https://portal.genego.com/) to test for enrichment in Pathway Maps, Process Networks, Diseases (by Biomarkers) and GO Processes (Table S24-27). MetaCore determines the significance of enrichment using a hypergeometric model based on the size of the intersection between the uploaded dataset and the genes/proteins in a given pathway.

## ACKNOWLEDGEMENTS

Olivia Newton-John Cancer Research Institute acknowledges the support of Operational Infrastructure Program of Victorian Government.

## AUTHOR CONTRIBUTIONS

NP and DD conceived the study and wrote the manuscript. SH-K and RPR completed the bulk of the experimental work and contributed to the preparation of the manuscript. LC, XL and LCBM contributed to the experimental work. AJL and RCR prepared the HDACi and DPF prepared the HDAC inhibitors and critically revised the manuscript. RLA provided intellectual input and contributed to the review and editing of the manuscript. All authors reviewed the manuscript.

## Abbreviations

BBB: Blood-brain barrier
BCM: Brain conditioned medium
BrCa BSM: Breast Cancer Brain-Specific Metastasis
CTC: Circulating tumour cell
DAPI,: 4′,6-diamidino-2-phenylindole
ER: Oestrogen receptor
FBS: foetal bovine serum
FACS: fluorescence-activated cell sorting
GFAP: Glial fibrillary acidic protein
H&E: Hematoxylin and eosin
HDACi: Histone deacetylase inhibitor
HER2: Human epidermal growth factor receptor 2
PBS: Phosphate-buffered saline
PR: Progesterone receptor
RTB: Relative tumour burden
SFM: Serum-free medium
SRB: Sulforhodamine B
TNBC: Triple Negative Breast cancer

